# MicroRNAs buffer genetic variation at specific temperatures during embryonic development

**DOI:** 10.1101/444810

**Authors:** Christopher Amourda, Jeronica Chong, Timothy E. Saunders

## Abstract

Successful embryogenesis requires the coordination of developmental events. Perturbations, such as environmental changes, must be buffered to ensure robust development. However, how such buffering occurs is currently unknown in most developmental systems. Here, we demonstrate that seven miRNAs are differentially expressed during *Drosophila* embryogenesis at varying temperatures within natural physiological ranges. Lack of miR-3-309, -31a, -310c, -980 or -984c causes developmental delays specifically at a given temperature. Detailed analysis on miR-310c and -984c shows that their targets are typically mis-expressed in mutant backgrounds, with phenotypes more pronounced at temperatures where miRNAs show highest expression in wild-type embryos. Our results show that phenotypes may arise at specific temperatures while remaining silent at others, even within typical temperature ranges. Our work uncovers that miRNAs mask genetic variation at specific temperatures to increase embryonic robustness, highlighting another layer of complexity in miRNA expression.

## Introduction

Embryonic development is a succession of stereotypical events occurring in a timely manner. As a consequence, the ability to respond to “perturbing” cues, such as environmental changes, is a key feature to all organisms on Earth (Ashton et al., 2017; Kuntz and Eisen, 2014). The progression of a particular event needs to be coordinated to the timescale of development across the whole embryo. For example, fruit fly embryonic developmental time varies greatly with temperature, yet developmental events remain very precise and occur at similar relative times during the course of embryogenesis (Chong et al., 2018; Kuntz and Eisen, 2014). In zebrafish embryos, the period of the segmentation clock changes by a factor of three between 20°C and 30°C. However, the length of embryos remains constant (Schröter et al., 2008). The response to temperature is presumably unequal between biological processes, hinting that mechanisms must be in place to ensure robust embryonic development in the face of perturbations. A recent study revealed that the β-tubulin97EF is upregulated at 14°C in the gut and in haemocytes and is essential to stabilise microtubules at low temperature (Myachina et al., 2017). This illustrates the need to differentially regulate key cellular components to cope with abnormal temperatures.

While specific cases have been characterized, general principles still need to be unravelled (Ebisuya and Briscoe, 2018). Molecularly, microRNAs (miRNAs), which appeared concomitantly with the rise of multicellular organisms (Peterson et al., 2009), have been shown to ensure robustness of complex developmental processes (Alberti and Cochella, 2017; Hornstein and Shomron, 2006; Pelaez and Carthew, 2012; Reinhart et al., 2000; Schmiedel et al., 2015; Shomron, 2010). Since their discovery over two decades ago, numerous studies have established that their expression is tightly linked to environmental conditions (Leung and Sharp, 2010; Li et al., 2009). MiRNAs, therefore, play an essential role in maintaining homeostasis against internal and external perturbations. However, these effects are typically subtle (many miRNA homozygous mutants are viable), and it has proven challenging to infer miRNA function during development.

Experiments that have probed miRNA response to environmental cues have typically considered extreme or fluctuating conditions (Cassidy et al., 2013; Li et al., 2009). Yet, given the robustness of development at normal temperatures, we reasoned that it is likely that miRNAs could well be buffering variability within more physiological conditions. The fruit fly, *Drosophila melanogaster*, provides an excellent model to study such subtle variations as the whole organism can be easily raised and maintained at a specific temperature. Its developmental processes are extremely robust, allowing to score for phenotypes from embryonic to adult stages. Specifically, we probed the expression profiles of miRNAs at the lower, mid and upper ends of typical thermal conditions. Importantly, our study was performed to minimise environmental variability at each chosen temperature (18°C, 21°C, 25°C). We then identified miRNAs with varying expression profiles at different temperatures and carefully examined the mutant phenotypes at different temperatures. We identified temperature-specific phenotypes, suggesting that these miRNAs are playing a role in buffering variability to temperature. Though our work is in *Drosophila,* miRNAs are generally conserved across taxa in terms of targets (Chandra et al., 2017). Our study highlights miRNA classes that are differentially expressed at different temperatures, and the subsequent cellular effects of such expression variation with possible relevance to higher organisms, wherein such a study is currently largely unfeasible.

## Results and Discussion

### Identification of miRNAs differentially regulated within physiological temperature scales

To probe the miRNA expression under physiological conditions, we collected OregonR (OreR, or wild-type) embryos at the blastoderm stage of development and allowed them to develop until the onset of germ band retraction at 18, 21 or 25 (±0.5) °C. These temperatures are those classically used to maintain laboratory flies and are considered non-stressful to *Drosophila*. We extracted the total RNA from these embryos and performed an Affymetrix GeneChip miRNA array (Fig. 1A). We obtained a list of 22 miRNAs showing at least a two-fold expression level difference between two temperatures (Figs. 1B, 1C and S1). MiRNAs originating from a single cluster were found to display similar expression trends. For example, miR-3, -4, -6 and -309, from the miR-309 cluster, have highest expression levels at 18°C. Similarly, members of the miR-310 cluster show higher expression at 18°C compared to 21 and 25°C. Furthermore, the expression level of miR-7 is increased at 18°C compared to higher temperatures. This finding agrees with previous work showing that miR-7 is essential to maintain robust developmental programs in the presence of temperature fluctuations (Li et al., 2009).

**Figure 1:**
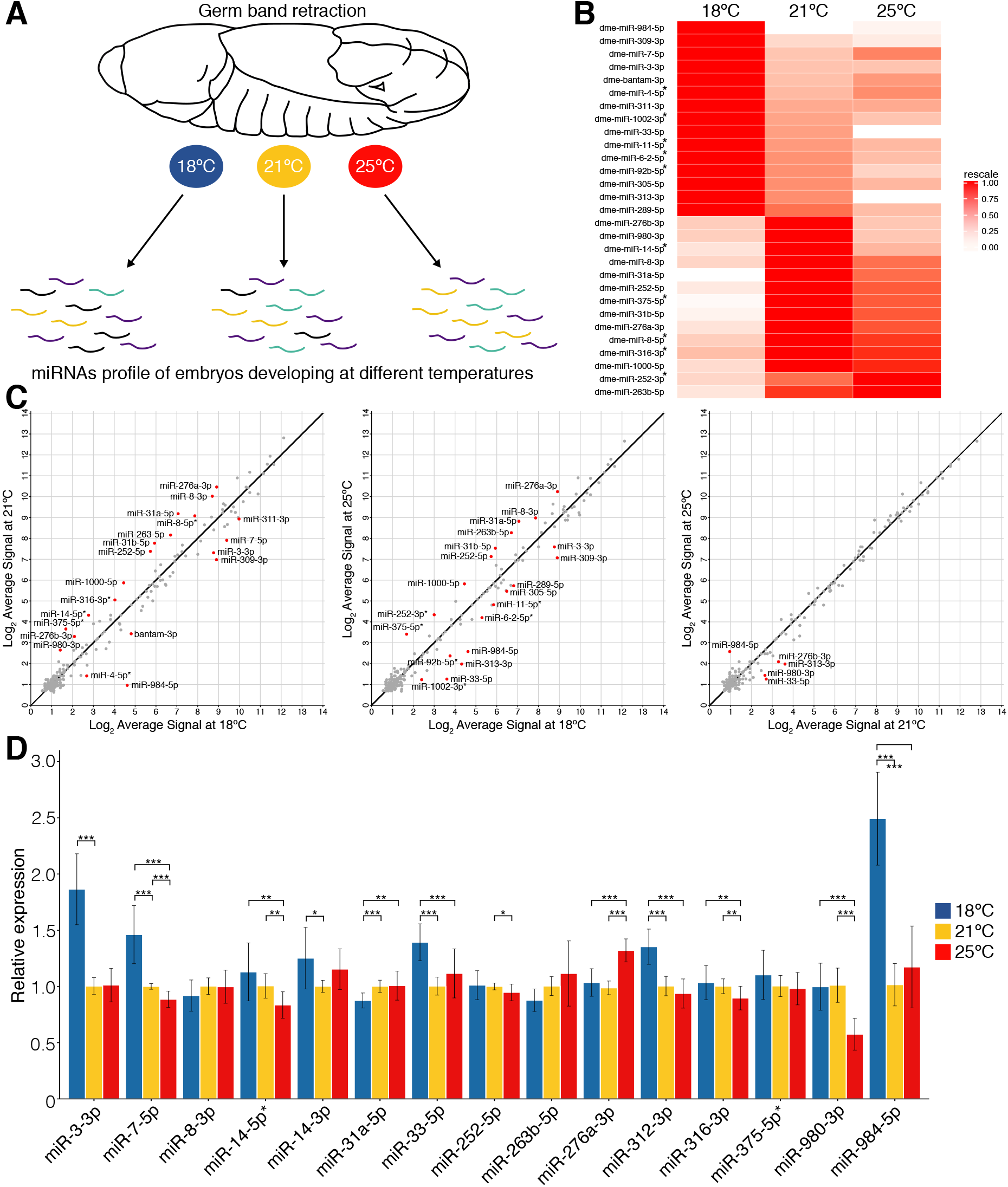
miRNA expression profiles vary with small temperature shifts. **(A**) Schematic of experimental procedures. 50 blastoderm embryos were allowed to develop until germ band retraction under precise temperature control (18, 21 and 25 ±0.5 °C). MiRNA profiles were obtained by Affymetrix GeneChip miRNA array. (**B**) Heat map of miRNAs showing more than 2-fold difference between different temperatures. Data normalised to the highest expression value for each miRNA. Relative expression level was color-coded to range from white (low levels of miRNA expression) to red (higher level of expression). Star species are denoted with a * after miRNA name. (**C**) Scatter plot comparing the expression level (log2 scale) between 18-21°C (right), 18-25°C (middle) and 21-25°C (left). miRNAs showing an expression difference equal to or greater than 2-fold are shown as red dots. (**D**) miRNA RT-qPCR expression level relative to the 21°C expression level for each miRNA. All values are means ± SD (*P<0.05, **P<0.01, ***P<0.001, n = at least 9 for each miRNA).

The formation of miRNAs give rise to a predominant guide strand and a subsidiary (complementary) star strand. MiRNA star species, although less abundant than their counterpart, can be present at physiologically relevant levels and show inhibitory effects similar to predominant miRNAs (Okamura et al., 2008). Our microarray results reveal that there can be temperature-specific upregulation of miRNA star strands (* in Fig. 1B), while the predominant strand remains unchanged at different temperatures. This suggests that miRNA star species could be involved in previously unknown temperature-dependent regulation.

The Affymetrix Chip microarray can often lead to false positives, even with sufficient biological replicates. Therefore, we confirmed candidates from the microarray by miRNA RT-qPCR. We found that miR-3, -7, -14, -33, -310c and -984 have significantly higher expression at 18°C while only miR-276b exhibit highest expression at 25°C (Fig. 1D).

### miRNAs regulate the timing of development events in a temperature-specific manner

Several miRNAs highlighted in our analysis are dispensable for viability and fertility under typical environmental conditions. As a consequence, their specific function has often remained elusive (Chen et al., 2014), particularly as most studies are carried out at 25°C. We reasoned that the timing of developmental landmarks, typically precise (Fig. S2 and Chong et al., 2018), may be altered at a given temperature in miRNA null background. We tested this hypothesis by scoring the time to reach specific developmental landmarks during embryonic development (Chong et al., 2018; Kuntz and Eisen, 2014) of miRNA null mutants in comparison to OreR embryos (Fig. S3). Among the list of candidates, miR-3-309, -31a, -310c, -980 and -984c mutants are homozygous viable and fertile. Mutant embryos were imaged alongside OreR embryos (Fig. 2A and Video 1). Under these imaging conditions, 90 to 95% of OreR embryos hatched as larvae at both 18 and 25°C, a rate similar to that observed in miR-31a and -980 mutants. In contrast, miR-3-309 mutants displayed a decrease to 75% of survival at 25°C. MiR-310c mutants have reduced survival at both 18 and 25°C, respectively to 80 and 65%. The miR-984c mutant survival rate is slightly inferior at 18°C (77%) compared with 25°C (85%) (Fig. 2B). Whether there are biologically significant differences in survivability at different temperatures remains to be tested.

**Figure 2:**
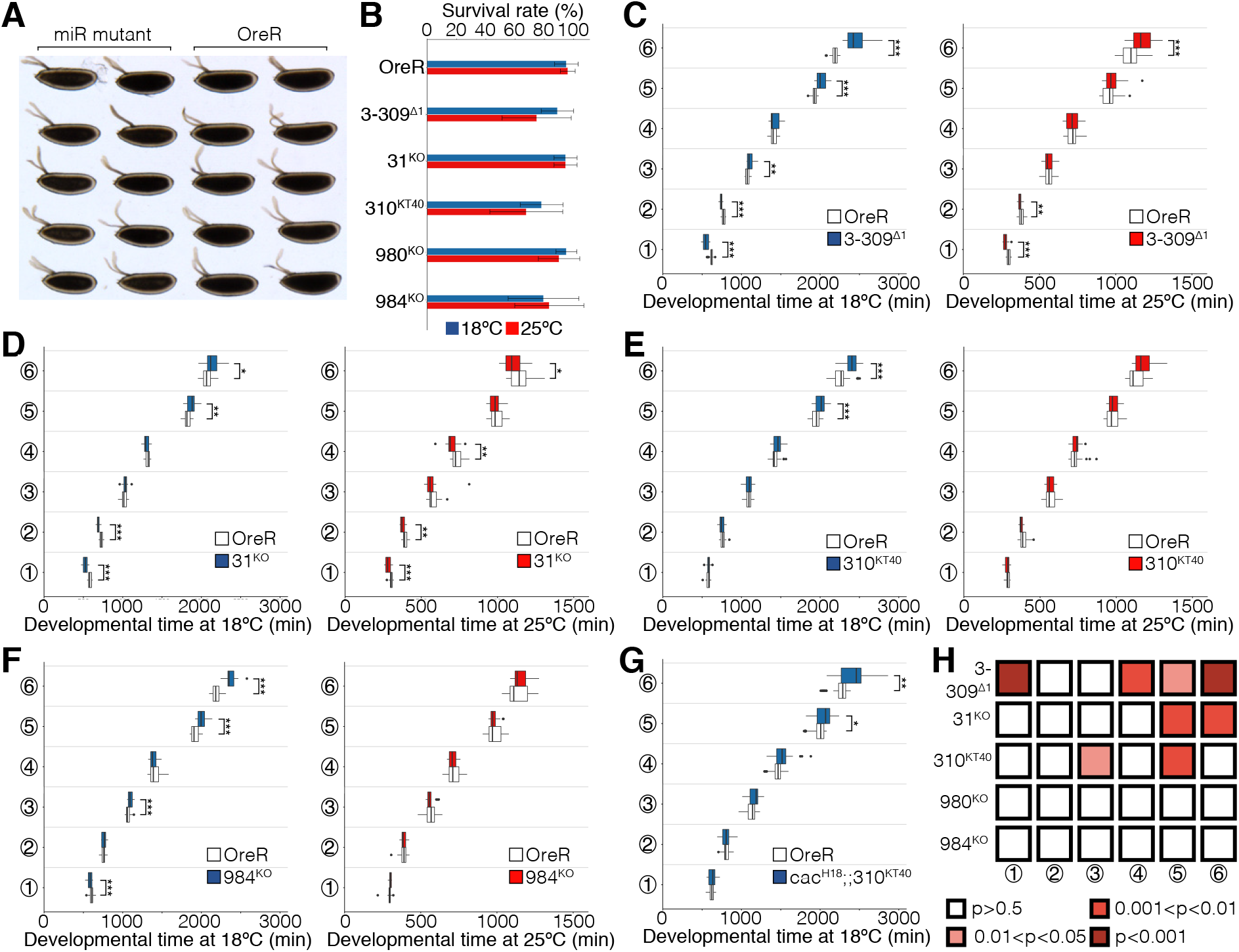
miRNA mutant analysis reveals developmental delay at specific temperatures. (**A**) Embryos laid on an apple juice agar plate. Ten OreR (right side) and 10 miRNA mutants (left side) embryos are imaged simultaneously at 18 or 25(±0.5) °C. (**B**) Survival rate measured in miRNA mutant embryos raised at 18 or 25°C. (**C-G**) Boxplot showing the developmental time scored in OreR and mutant embryos raised at 18 (blue) or 25 (red)°C. Landmarks are 1-Germ band retraction, 2-Head involution, 3-Midgut broadening, 4-Muscle contraction, 5-Trachae fill, 6-Larvea hatching. Mutant embryos are miR-3-309^∆1^ (**C**), miR-31^KO^ (**D**), miR-310c^KT40^ (**E**), miR-984^KO^ (**F**) and cac^H18^; miR-310c^KT40^ (**G**). (**H**) Robustness analysis at 18°C, comparing standard deviation of temporal trajectories between mutants and OreR embryos, see Methods. P-values presented describe whether there was a statistically significant increase in the temporal variability in mutant conditions. Between 15 to 20 embryos were imaged and analysed for each miRNA mutant. Statistical significance is shown as *P<0.05, **P<0.01, ***P<0.001.

We next focused on the variation between specific developmental landmarks (Methods – Experimental reproducibility for details); *i.e.* we predicted that the miRNAs are playing a role in temporal coordination of embryonic development. MiR-3-309^∆1^ (Bushati et al., 2008) embryos exhibited both a general developmental delay and an increase in temporal variability (see below) at both temperatures compared to OreR, with larval hatching delayed by 336 and 70 min at 18°C and 25°C respectively (Fig. 2C). Analysis of the time difference between our recorded developmental landmarks in miR-3-309^∆1^ embryos revealed that the sequence of developmental events is most dramatically affected at 18°C, with the onset of head involution, midgut broadening and trachea fill being significantly delayed by 40, 62 and 70 mins, respectively (Fig. S4A). MiR-31a^KO^ (Chen et al., 2014) embryos showed a similar trend, with embryogenesis affected at both 18 and 25°C (Fig. 2D). However, miR-31a has a larger effect later in development, with a delay of 60min at 18°C but faster development at 25°C (46 min) (Fig. S4B). MiR-310c^KT40^ (Tsurudome et al., 2010) embryos hatched significantly later at 18°C (120min) but not at 25°C (Fig. 2E) when compared to wildtype embryos. In particular, the onset of muscle contraction was delayed by 40min, explaining the hatching delay (Fig. S4C). MiR-984c^KO^ (Chen et al., 2014) embryos displayed a developmental delay (216min) solely at 18°C (Fig. 2F) with the onset of head involution, midgut broadening and trachea fill being significantly delayed by 26, 22 and 98min, respectively (Fig. S4D), while the onset of muscle contraction was quickened by 48min. This pattern is reminiscent to that observed in miR-310c^KT40^ embryos. Whilst overall embryogenesis appeared unaffected in miR-980^KO^ (Chen et al., 2014) embryos, the developmental choreography was affected most at 18°C (Fig. S4E).

To validate that the timing phenotype in miR-310c^KT40^ results from the lack of miRNAs, we attempted to rescue miR-310c^KT40^ as its functional targets are well characterised. In the absence of miR-310c, the levels of Khc-73 increases and, as a result, Bruchpilot (Brp) shows high accumulation at the synapse. Brp facilitates the localisation of Cacophony (Cac) in presynaptic zones, triggering neurotransmitter release. MiR-310c^KT40^ are rescued in Cac heterozygous mutants (Tsurudome et al., 2010). Corroborating these results, we managed to partially rescue the timing delay observed in miR-310c^KT40^ by reducing Cac levels to 50% (Fig. 2G). Further, we attempted to fully rescue miR-310c^KT40^ by exogenous insertion of miR-310c genomic locus onto the 3^rd^ chromosome (Fig. S5A). Unfortunately, the genomic region cloned failed to rescue the timing phenotype (Fig. S5B) and, further, the miR-310c level remains low (~30%) in the rescue line (Fig. S5C). This indicates that the regulatory regions controlling the expression of miR-310c are more complex than initially thought. Nevertheless, these experiments allowed us to correlate the miRNA expression data to phenotypes, corroborating that classes of miRNAs play differing orles at a given temperature. The developmental landmarks affected can be pinpointed, allowing targeted future studies of the underlying mechanisms.

### miRNAs regulate developmental timing precision at specific temperatures

We next asked whether there was a change in the temporal robustness (variability between trajectories under the same conditions) with loss of specific miRNAs? We calculated the standard deviation in the temporal trajectories at each landmark for each genotype and compared these with the variation in OreR embryos recorded alongside. Uncertainty on our estimates of the standard deviation were calculated using bootstrapping, see Methods. At 18°C, we found that 3-309^Δ1^, 31a^KO^, and 310c^KT40^ embryos all showed greater temporal variability, particularly towards the end of embryogenesis, consistent with temporal defects accumulating during embryogenesis due to decreased coordination in the mutants (Fig. 2H). Since these effects are likely subtle, the cumulative effect only becomes apparent later in development. Conversely, at 25°C, there was no statistically significant loss of temporal robustness in these embryos toward the end of embryogenesis. Temporal robustness was largely unaffected in early embryogenesis in 980^KO^ and 984c^KO^ embryos (Fig. 2H). However, this does not discount a loss of robustness in other aspects of development in these embryos.

### MiR-984c and miR-310c targets show temperature-specific upregulation

To understand the underlying mechanisms to temperature sensitive temporal regulation of development by miRNAs, we focused our attention on miR-984c and -310c. Mutants of these both show a clear phenotype arising exclusively at 18°C. We reasoned that the expression level of miR-984c and -310c targets should differ between 18 and 25°C in the mutant backgrounds.

Since the roles of miR-984c are poorly understood, we used the predicted targets from the miRANDA (Betel et al., 2008) and the TargetScan (Kheradpour et al., 2007) fly databases and shortlisted 24 targets identified by both databases. We found that the level of *CG7220*, *Stem cell tumor* (*Stet*) and *Regeneration (Rgn*) mRNA is significantly higher (by 1.44±0.06-, 2.44±0.33- and 1.58±0.30-fold, respectively) at 18°C than in OreR or miR-984c^KO^ raised at 25°C. These results confirmed that miR-984c plays an important role at 18°C to maintain mRNA levels. Noticeably, the levels of *CG18625*, *Zelda*, *Wech* and *CG7987* mRNA are significantly lower (to 0.22±0.05-, 0.47±0.30-, 0.67±0.04- and 0.83±0.14-fold) at 18°C as compared to control embryos or to miR-984c^KO^ embryos raised at 25°C (Figs. 3A and S5). Among the list of potential targets, our results argue that *GC7720*, *Stet*, *Rgn* are functional targets of miR-984c as their level is higher in miR-984c^KO^. The low levels of *CG18625*, *Zelda*, *Wech* and *CG7987* indicates that they are not directly targeted by miR-984c and that their expression levels result from altered developmental processes. Further, the expression of Zelda - a DNA-binding protein controlling chromatin accessibility, transcription factor binding and gene expression (Schulz et al., 2007) - is of particular importance. Some tissues may be unable to undergo chromatin rearrangement to allow the embryo to trigger subsequent developmental events. This reduction in Zelda could possibly be due to an earlier defect in temporal coordination of developmental events resulting from the high level of *Stet*, which is involved in determining germ line cell fate (Guichard et al., 2000; Wasserman et al., 2000). Another interesting result here is the identification of previously uncharacterized genes in potentially ensuring robust development (e.g. CG7220, CG7987, and CG18625).

**Figure 3:**
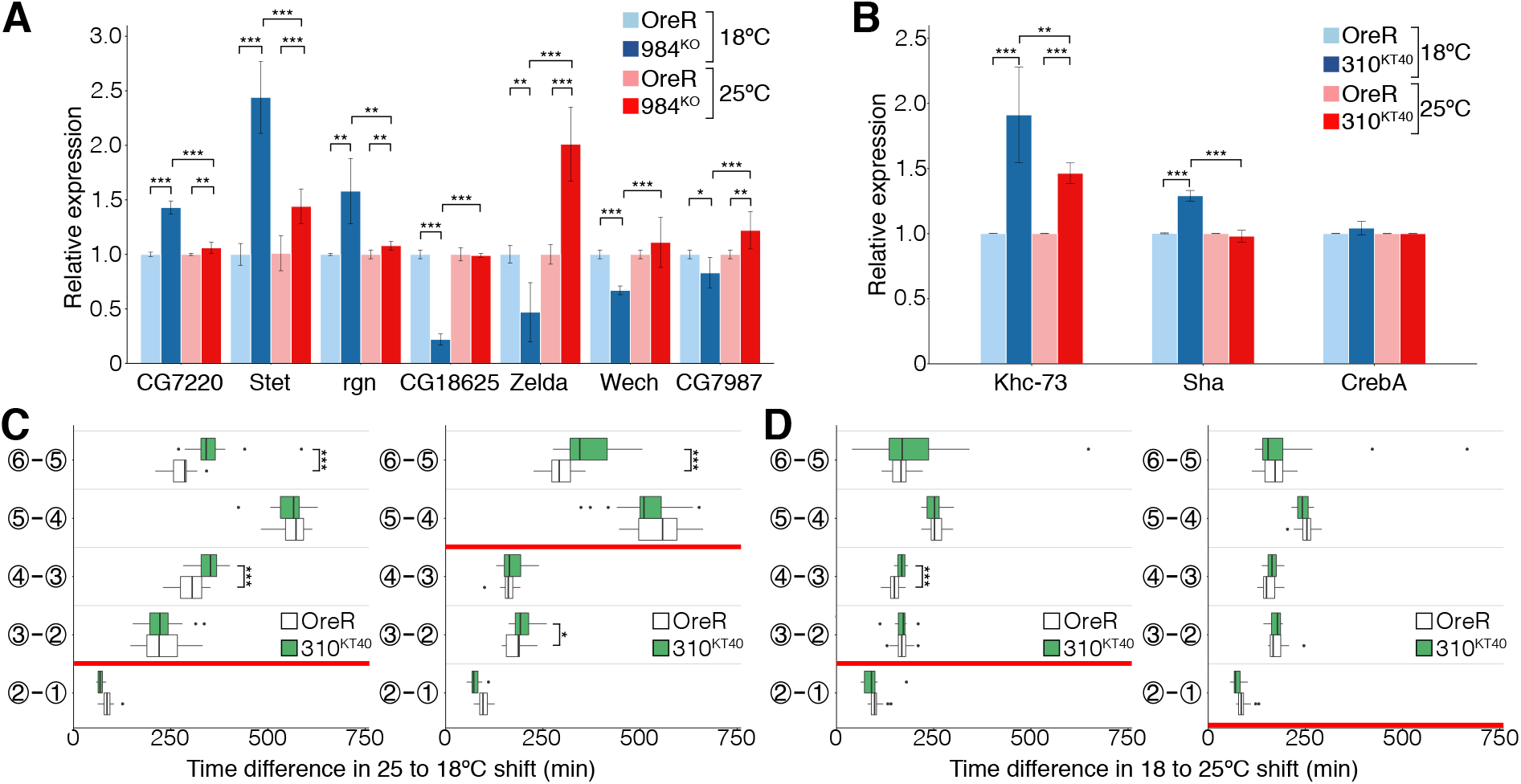
miR-984c and miR-310c targets show highest variation at 18°C. (**A**) Histogram of miR-984 targets expression in miR-984^KO^ background at 18 (dark blue) and 25 (dark red) °C relative to OreR embryos raised at 18 (light blue) and 25 (light red) °C. (**B**) Histogram of miR-310c validated targets in miR-310c^KT40^ background at 18 (dark blue) and 25 (dark red) °C relative to OreR embryos raised at 18 (light blue) and 25 (light red) °C. For both histograms, values are means ± SD (*P<0.05, **P<0.01, ***P<0.001, n= 9 for each miRNA). (**C-D**) Boxplot showing the time difference between two developmental landmarks scored in OreR and miR-310^KT40^ embryos. Red bars indicate the time at the temperature was shifted from 25 to 18°C (**C**) and from 18 to 25°C (**D**). In (**D)** right panel, the temperature has been shifted just before germ band retraction. Encircled number indicates the developmental stage described in legend to Fig. 2 and Fig. S2A. Between 15 to 20 embryos were imaged and analysed for each miRNA mutant. Statistical significance is shown as *P<0.05, **P<0.01, ***P<0.001.

### Lack of miR-310c results in a temperature-specific lack of embryonic robustness

Roles of miR-310c are generally better understood (Çiçek et al., 2016; Li et al., 2017; Pancratov et al., 2013; Tsurudome et al., 2010). We, therefore, focused our analysis on validated targets and highest hits from target prediction tools. We found that expression of Kinesin heavy chain (Khc)-73 shows a 1.91±0.36-fold increase at 18°C, significantly larger than the increase seen at 25°C (1.46±0.08-fold). Shavenoid (Sha) is mildly overexpressed (1.29±0.04-fold) at 18°C as compared to embryos raised at 25°C. Last, the expression of the CrebA is not affected in miR-310c^KT40^ embryos (Fig. 3B). MiR-310c has been shown to control normal synaptic transmission at the larval neuromuscular junction by modulating the Khc-73 level (Tsurudome et al., 2010). Hence, we hypothesised that miR-310c is critical to central nervous system (CNS) morphogenesis and ensures robust timing of muscle contractions at lower temperatures. To corroborate these results, we sought to rescue the developmental delay by switching the temperature from 18 to 25°C; the delay can be rescued by switching temperature before head involution but not later (Fig. 3C). The opposite experiment, switching the temperature from 25 to 18°C, showed that the delay only occurs when the temperature is switched before the muscle contraction stage (Fig. 3D). These results suggest that the mis-expression of Khc-73 accounts for the timing delay observed in miR-310c^KT40^ embryos at 18°C.

To better understand the phenotypes resulting from the over-expression of Khc-73, we performed live-imaging of embryos expressing sGMCA in a miR-310c^KT40^ background (Video 2). As predicted, we found that the developmental delay occurs at the onset of muscle contraction with the full CNS condensation taking on average 102±86mins more than control embryos. The kymograph trace of the CNS persists longer in miR-310c^KT40^ embryos raised at 18°C as compared to control embryos and embryos maintained at 25°C (Fig. 4A). This observation suggests that the ability to twitch muscles is more dramatically affected at 18°C. Altogether, these results are in accordance with Tsurudome *et al.* (Tsurudome et al., 2010) who showed that the de-repression of Khc-73 in miR-310c^KT40^ causes an increase in Ca^2+^ influx presynaptically, resulting in an increase in quantal content. Therefore, miR-310c plays a critical role in regulating CNS maturation at lower temperatures.

**Figure 4:**
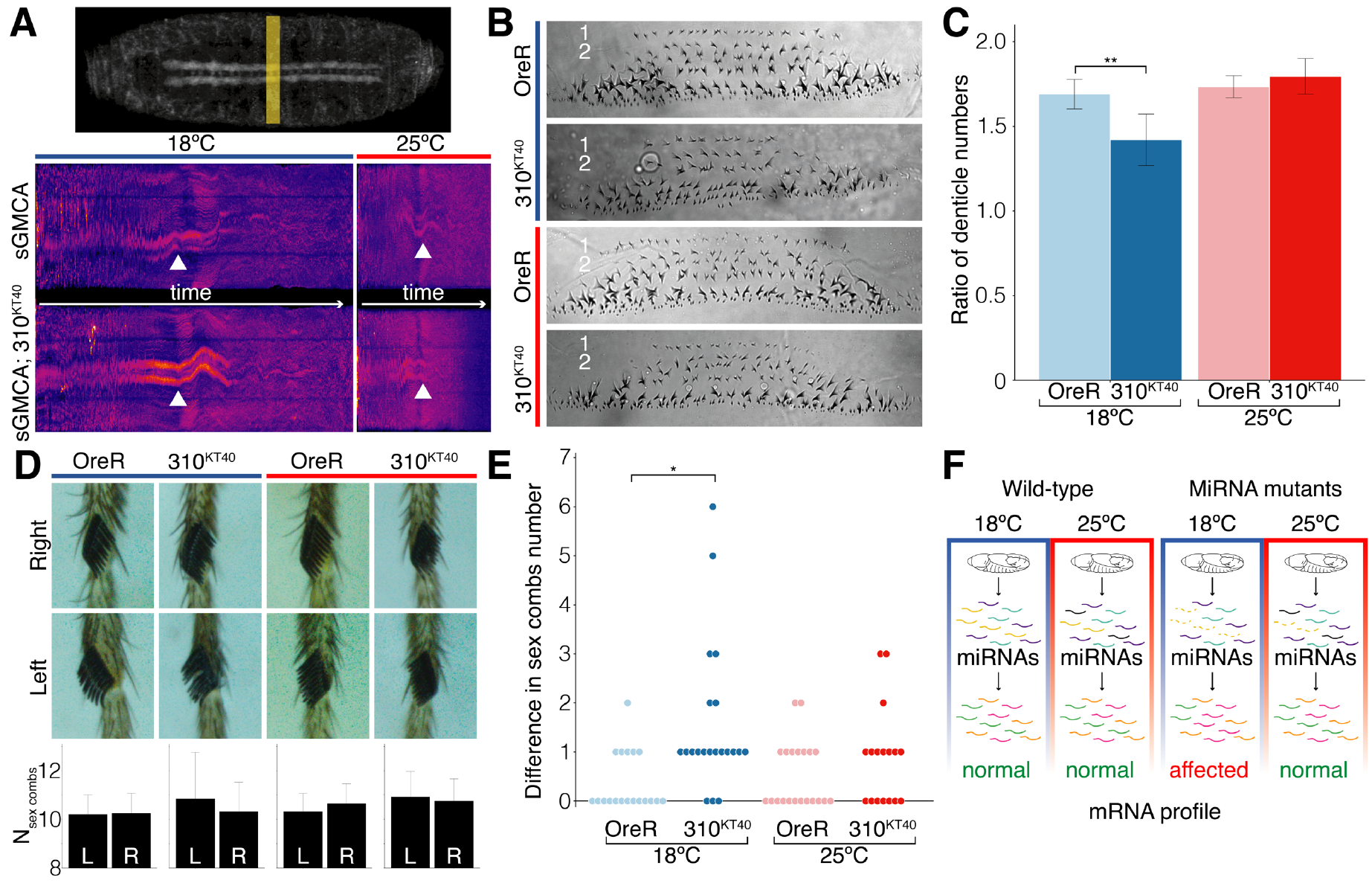
miR-310c^KT40^ ensures robust development at low temperatures. (**A**) Yellow box in the top panel indicate the area where kymographs were performed. Bottom panels shows Kymographs for sGMCA (top) and sGMCA; miR-310c^KT40^ (bottom) embryos imaged at 18 (left panels) and 25 (right panels) °C. White arrowheads indicate the position of the CNS. (**B**) Denticle pattern observed in OreR and miR-310c^KT40^ embryos raised at 18 (blue) or 25 (red) °C. (**C**) Ratio of the number of denticles between row 1 and 2. Data are presented as means ± SD (**P<0.01, n = 5 larvae per condition). (**D**) Representative sex combs in OreR and miR-310c^KT40^ adult males raised at 18 (blue) or 25 (red) °C. Bottom panel show the average number (± SD) of sex combs counted on the left and right leg in each condition. (**E**) Dot plot of the difference in sex combs number between the left and right legs for OreR raised 18 (light blue) or 25 (light red) °C and miR-310cKT40 raised at 18 (dark blue) or 25 (dark red) °C (*P<0.05, n = 17-20 pair of legs per condition). (**F**) Schematic representation of an example of miRNAs regulation at different temperatures.

### miR-310c has temperature-specific roles in regulating morphogenesis

The above reported fold changes in miRNA expression at different temperatures are generally small. Therefore, we next investigated whether such alterations have a morphological effect during development. We focused on miR-310c, and examined both embryonic and adult structures at different temperatures in a miR-310c mutant background. We calculated the number of larval denticles in miR-310c^KT40^ embryos raised at 18 and 25°C. For unambiguous calculation, we evaluated the number of denticles in row 1 and 2 (Fig. 4B). The ratio row1/row2 shows that wild-type and miR-310c^KT40^ embryos have similar denticle patterns at 25°C. Raising miR-310c^KT40^ embryos at 18°C causes variable denticle patterns with often the presence of ectopic denticles (Fig. S7). The ratio row1/row2 is reduced to 1.42±0.15, compared with 1.71±0.07 at 25°C (Fig. 4C). These results are consistent with mildly higher levels of Sha in miR-310c^KT40^ raised at 18°C. While the effects of Sha over-expression are unknown, its mutants show strong denticle defects (Chanut-Delalande et al., 2006). Lastly, we tested whether miR-310c temperature dependence is restricted to embryogenesis or rather a general feature of this cluster by looking at morphological abnormalities in miR-310c^KT40^ adults raised at 18 and 25°C for several generations. The number of leg sex combs shows an increase from 10.4±0.2 to 10.7±0.3 in the absence of miR-310c at both temperatures. While the change in mean number is small, the variation in sex comb number is doubled in miR-310c^KT40^ raised at 18°C (Fig. 4D), and left-right asymmetry in sex comb number showed a significant increase (Fig. 4E). Hence, miR-310c has a general role in ensuring robust development at lower temperatures.

Taken together, our screen has uncovered the existence of physiologically relevant differential miRNA expression within the range of natural environmental conditions. We demonstrate that miRNAs act to maintain robust mRNA levels under relatively small temperature changes (Fig 4F), suggesting that temperature regulation during *Drosophila* development – via miRNAs – is much more finely tuned than previously believed. Crucially, our results also reveal that phenotypes may arise at specific temperatures while remaining silent at others, even within typical temperature ranges. As the global climate changes plants and animals will be exposed to increasing (and increasingly variable) temperatures. Our work reveals that understanding how life will adapt to such changes (though seemingly small at around 1-2°C over the next century) requires a more complete comprehension of the role of miRNAs, which appear to play a central role in regulating small temperature variations.

Having established that temperature mediates differential miRNA expression, understanding the mechanisms that control this thermal-dependence will pose an interesting challenge. For instance, the posttranscriptional addition of an uridine or an adenosine at the 3’ end of miRNAs has been shown to control miRNA decay, processing and activity (Burroughs et al., 2010; Katoh et al., 2009; Lee et al., 2014). Deep sequencing analyses of embryos raised at different temperatures may reveal distinctive 3’ capping. Alternatively, the chromatin organisation may change at different temperatures. Indeed, a study in plants established that a histone variant (H2A.Z) is differentially localised on the genome of *Arabidopsis thaliana* and that this localisation varies with temperatures (Kumar and Wigge, 2010). This allows the expression of certain genes at particular temperatures while at the same time repressing other genes. Such histone variants are also present in animals, and reportedly show specific localisation on the genome (Leach et al., 2000). An attractive possibility is that temperature changes the distribution of the histone variant on the genome, which in turns regulates the transcriptome. Overall, our work demonstrates a surprisingly rich diversity of behaviour *in vivo* with relatively small temperature changes and paves the way to future studies understanding cell responses to temperature changes.

## Acknowledgments

We thank Pejmun Haghighi and Stephen Cohen for sharing precious reagents. We acknowledge Stephen Cohen, Sherry Aw, Katsutomo Okamura, Nicholas Tolwinski, Yusuke Toyama and all Saunders’ lab members for fruitful discussions. This work was supported by the National Research Foundation Singapore under an NRF Fellowship to T.E.S. (NRF2012NRF-NRFF001-094).

## Author Contributions

C.A. and T.E.S. designed the study. C.A. and J.C. performed the experiments. C.A. analysed the data. C.A. and T.E.S. wrote the manuscript. All authors participated in the final version of the manuscript.

The authors declare no competing financial interest.

